# Spatiotemporal differences of GABAergic polarization and shunting during dendritic integration

**DOI:** 10.1101/2025.04.05.647352

**Authors:** Yulia Dembitskaya, Artem Kirsanov, Yu-Wei Wu, Alexey Brazhe, Alexey Semyanov

**Affiliations:** Shemyakin-Ovchinnikov Institute of Bioorganic Chemistry, Russian Academy of Sciences, Moscow, Russia 117997; Moscow State University, Faculty of Biology, Moscow, Russia; Institute of Molecular Biology, Academia Sinica, NanKang, Taipei, Taiwan; Department of Physiology, Jiaxing University College of Medicine, Zhejiang, 314033 China

**Keywords:** inhibition, excitation, dendritic integration, depolarizing GABA, hyperpolarizing GABA, shunting

## Abstract

In the adult brain, GABA exerts either depolarizing or hyperpolarizing effects on neuronal membranes, depending on neuron type, subcellular location, and neuronal activity. Depolarizing GABA typically inhibits neurons through shunting, which is characterized by increased membrane conductance upon GABA_A_ receptor activation; however, it can also excite neurons by recruiting voltage-dependent conductances. The net influence of these opposing actions of depolarizing GABA on glutamatergic synaptic inputs remains incompletely understood. Here, we examined the spatiotemporal characteristics of membrane polarization and shunting mediated by GABA_A_ receptors and assessed their functional impact on the integration of GABAergic and glutamatergic inputs along dendrites. Using whole-cell current-clamp recordings in CA1 pyramidal neurons and dentate gyrus granule cells (GCs) from rat hippocampal slices, we mimicked GABAergic and glutamatergic inputs with local GABA puffs and glutamate spot-uncaging, respectively. A mathematical model further quantified the relative effects of local shunting and polarization. Depolarizing GABAergic postsynaptic responses (GPSRs) exhibited biphasic actions, exerting inhibitory effects at the synapse through shunting, and excitatory effects distally, where depolarization predominated. The excitatory component also persisted longer than the shunting inhibition. In contrast, hyperpolarizing GPSRs remained consistently inhibitory across both spatial and temporal dimensions. These findings highlight the complex spatiotemporal interplay between shunting and membrane polarization mediated by GABAergic inputs, providing new insights into dendritic computation and neuronal network dynamics.

## Introduction

Although GABA is the principal inhibitory neurotransmitter in the brain, its effects on neuronal excitability and synaptic integration are highly nuanced. The spatiotemporal integration of excitatory and inhibitory inputs in dendrites critically determines neuronal action potential (AP) output and influences information processing efficiency in neural circuits (Baufreton et al., 2005; Lee et al., 2022).

GABA acts on the neuronal membrane primarily via ionotropic GABA_A_ receptors, which are permeable to Cl^−^ and HCO_3_^−^ (Takeuchi and Takeuchi, 1967; Farrant and Kaila, 2007). In adulthood, the intracellular Cl^−^ concentration decreases, mainly due to enhanced expression of K/Cl^−^ co-transporter 2 (KCC2)(Perkins and Wong, 1997; Karten et al., 2006; Valeeva et al., 2016), resulting in the hyperpolarizing effect of GABA. In adulthood, the intracellular Cl^−^ concentration decreases, primarily due to enhanced expression of K/Cl^−^ co-transporter 2 (KCC2) (Payne et al., 1996; Rivera et al., 1999), resulting in the hyperpolarizing effect of GABA. Nevertheless, GABA can evoke depolarizing or hyperpolarizing responses depending on neuronal subtype, cellular compartment, and physiological state (Gulledge and Stuart, 2003; Pugh and Jahr, 2011; Chiang et al., 2012; Haam et al., 2012). Furthermore, hyperpolarizing GABAergic responses may transition to depolarizing under conditions such as changes in circadian phase (Pracucci et al., 2023) or pathological states (Jaenisch et al., 2010; Weilinger et al., 2022; Kim and Martina, 2024; McArdle et al., 2024; McMoneagle et al., 2024; Zhu et al., 2024).

Hyperpolarizing GABA typically suppresses neuronal firing by driving the membrane potential away from the AP threshold. However, it can also activate hyperpolarization-activated (*h*) channels, leading to membrane depolarization due to their mixed cationic conductance (Pavlov et al., 2011). Activation of *h*-channels can elicit rebound spikes, where APs follow inhibitory postsynaptic potentials (IPSPs), contributing to neuronal network synchronization (Kranig et al., 2013). Additionally, di-synaptic IPSP activation of *h*-channels shapes monosynaptic excitatory postsynaptic potentials (EPSPs) in CA1 pyramidal neurons (George et al., 2009), thus defining the temporal integration window for EPSPs.

Depolarizing GABAergic responses are documented in various adult neuronal populations, including hippocampal interneurons (Otsu et al., 2020), dentate gyrus granule cells (GCs) (Karten et al., 2006; Chiang et al., 2012), and cerebellar GCs (Dave and Bordey, 2009; Pugh and Jahr, 2011). Although depolarizing responses typically remain subthreshold, with the GABA current reversal potential (E_GABA_) below AP threshold, they can amplify membrane potential fluctuations caused by spontaneous Na^+^ channel openings, ultimately driving the membrane potential to AP threshold (Song et al., 2011; Seong et al., 2014).

In addition to membrane polarization, GABA_A_ receptor activation increases membrane conductance, causing shunting of EPSPs, thus reducing their amplitude and likelihood of reaching AP threshold (Chiang et al., 2012). This shunting either enhances the inhibitory effects of hyperpolarizing GABA or counters the excitatory influence of depolarizing GABA. Intriguingly, the excitatory action of depolarizing GABA critically depends on the level of GABA_A_ receptor-mediated conductance: low conductance permits excitation, whereas high conductance predominantly results in inhibition through shunting (Song et al., 2011; Chiang et al., 2012; Heigele et al., 2016).

Importantly, membrane conductance and polarization differ markedly in their spatiotemporal dynamics. Increased conductance due to GABA_A_ receptor activation is spatially localized at receptor sites (synapses) and returns rapidly to baseline levels. In contrast, polarization effects propagate along the membrane and persist longer due to membrane capacitance (Farrant and Kaila, 2007). Consequently, GABA_A_ receptor activation creates distinct spatiotemporal profiles of conductance and polarization across dendritic membranes (Lombardi et al., 2021a). These profiles are determined by membrane time and length constants, both dependent on membrane conductance.

Thus, the relative positioning and timing of GABAergic and glutamatergic synaptic activations dictate how GABAergic postsynaptic responses (GPSRs, characterized by both conductance and polarization) affect EPSPs. Several theoretical studies have investigated the spatiotemporal integration of GPSRs and glutamatergic EPSPs (Jean-Xavier et al., 2007; Hao et al., 2009; Gidon and Segev, 2012; Li et al., 2014; Paulus and Rothwell, 2016; Lombardi et al., 2021a). Computational modeling indicates that even mildly depolarizing GPSRs may switch from inhibitory to excitatory actions under specific conditions (Branchereau et al., 2016; Lombardi et al., 2021a). In this study, we present the first direct experimental evidence demonstrating that depolarizing GPSRs produce complex inhibitory and excitatory effects, varying spatially and temporally, as they integrate with glutamatergic EPSPs. Our results underscore the intricate and dynamic role of GABAergic signaling in dendritic integration and neuronal network function.

## Results

We performed whole-cell current-clamp recordings from CA1 pyramidal neurons in rat hippocampal slices (Fig. 1A). Neurons were loaded with Alexa Fluor 594 via the patch pipette, allowing morphological visualization using two-photon imaging (Fig. 1B). To mimic GPSRs, brief local applications of GABA (10 µM) were delivered via short pressure pulses (8 – 15 ms, 6 psi, “GABA puff”) at the midpoint of the apical dendrite (Supplementary Video 1). For glutamatergic responses, 400 µM 4-methoxy-7-nitroindolinyl (MNI)-glutamate was applied in the bath solution and locally photolyzed (“spot-uncaged”) at positions coincident with the GABA puff site (0 µm), or 50 and 100 µm toward (+50, +100 µm) or away from (−50, −100 µm) the soma. Uncaging produced excitatory postsynaptic potential (EPSP)-like responses recorded at the soma, here termed uncaging-induced EPSPs (uEPSPs). To examine temporal integration, we triggered uEPSPs at 0, 250, 500, 800, and 1000 ms delays after GPSR initiation.

**Figure 1.**
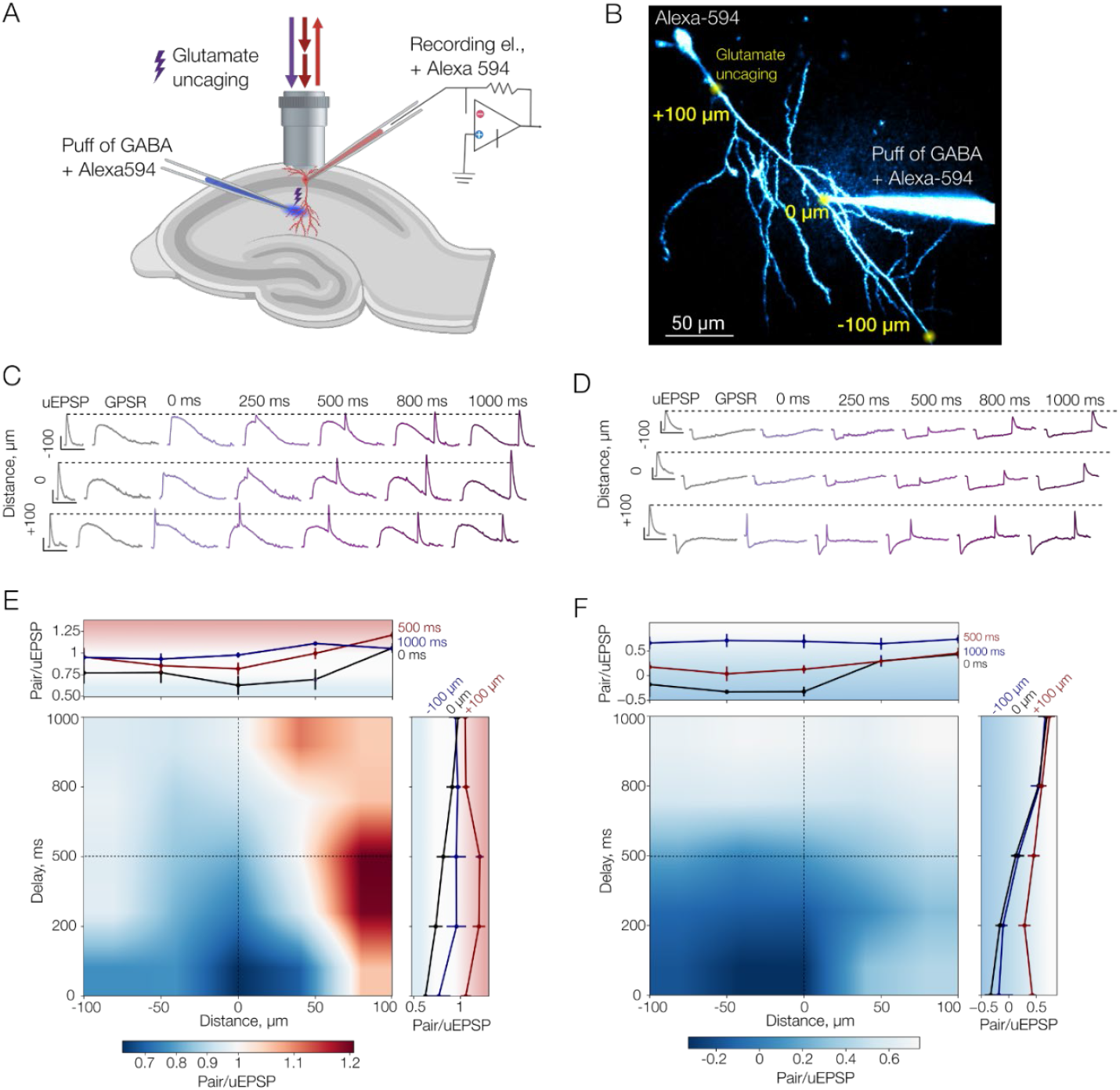
Depolarizing and hyperpolarizing effects of GABA in CA1 pyramidal neurons. **A**. Schematic of hippocampal slice, recording electrode, and puff pipette located in CA1 area. **B**. CA1 pyramidal neuron filled with Alexa Fluor 594 (50 μM) and puff pipette filled with Alexa Fluor 594 (15 μM). **C**. Sample traces of uEPSP, GPSR, and paired uEPSP + GPSR responses for depolarizing GABA. **D**. Sample traces of uEPSP, GPSR, and paired uEPSP + GPSR responses for hyperpolarizing GABA. **E**. Summary graph showing Pair/uEPSP ratio for different time delays and distances for depolarizing GABA. **F**. Summary graph showing Pair/uEPSP ratio for different time delays and distances for hyperpolarizing GABA. The data are presented as the mean ± SEM. For statistical data, see Supplementary tables 1-4.

Initially, we tested the effect of pairing uEPSPs with depolarizing GPSRs (E_GABA_ = −62.81 mV; resting membrane potential (RMP) = −67.07 mV; Fig. 1C). The paired response (uEPSP + GPSR, “Pair”) amplitude was normalized to uEPSP alone to quantify excitation (ratio > 1) or inhibition (ratio < 1). When uEPSPs were triggered proximally to the soma (+100 µm), the GPSR consistently exhibited biphasic excitatory effects (Fig. 1C, E; Supplementary Tables 1, 2). The strongest excitatory effect occurred with a 250 ms delay, decreasing gradually at longer intervals. Conversely, at the GABA puff site (0 µm), GPSRs exclusively exerted inhibition, even when the uEPSP was delayed relative to GPSR onset.

These results suggest that at the GABA application site, the inhibitory shunting effect predominates over excitation mediated by membrane depolarization. However, inhibition diminished more rapidly with distance compared to the depolarizing excitation. Temporally, shunting inhibition was constrained to periods of open GABA_A_ receptors, whereas depolarization persisted longer due to the membrane’s capacitive properties. Consequently, depolarization’s excitatory influence surpasses shunting inhibition both spatially and temporally. Nevertheless, the net effect depended on the relative dendritic positions of uEPSP initiation and GPSR sites. Distal dendritic uEPSPs (−100 µm) propagating toward the soma experienced *en route* inhibition when passing the GABAergic site. Although weaker and more transient than direct (co-localized) inhibition, this effect prevented the observation of excitation evident for proximally initiated uEPSPs (+100 µm). Thus, glutamatergic synapses near GABAergic inputs are inhibited by shunting, while proximal dendritic synapses relative to the soma are facilitated. Synapses distal to the soma encounter offsetting *en route* inhibition.

Next, we examined pairings of uEPSPs with hyperpolarizing GPSRs (E_GABA_ = −83.44 mV, RMP = −72.66 mV; Fig. 1D). Hyperpolarizing GABA consistently produced inhibitory effects, which depended on both temporal delay and spatial arrangement relative to the GPSR site (Fig. 1D, F; Supplementary Tables 3, 4). The strongest inhibition occurred at zero delay and co-localization (0 µm), but substantial inhibition was also evident for distal uEPSPs (−50, −100 µm), reflecting both membrane hyperpolarization and *en route* shunting. However, the inhibitory effects on proximal uEPSPs (+50, +100 µm) were comparatively weaker, primarily due to hyperpolarization without strong spatially restricted shunting.

These observations indicate that hyperpolarizing GABA prominently inhibits nearby glutamatergic synapses through localized shunting, while more distal synapses are inhibited by combined hyperpolarization and *en route* inhibition. Similar to depolarizing GABA, hyperpolarizing GABAergic inputs thus function as spatiotemporal gates modulating the somatic integration of proximally located glutamatergic synapses.

The above-described experiments were performed in CA1 pyramidal neurons, where GABA is typically hyperpolarizing in the adult brain (Scimemi et al., 2005). Nonetheless, a subcellular gradient of GABA reversal potentials, transitioning from hyperpolarizing at the soma to depolarizing in distal dendrites, has been proposed (Gulledge and Stuart, 2003), along with activity-dependent shifts toward depolarization (Payne et al., 1996; Rivera et al., 1999). Furthermore, depolarizing GABA actions have been documented in adult hippocampal interneurons (Song et al., 2011; Pavlov et al., 2014; Otsu et al., 2020) and dentate gyrus GSs (Karten et al., 2006; Chiang et al., 2012). Since GSs exhibit lower membrane conductance compared to CA1 pyramidal neurons (Kowalski et al., 2016), we hypothesized differences in spatiotemporal GPSR effects between these cell types.

To address this, we repeated the experiments using depolarizing GABA in GSs (Fig. 2A, B). Similar to pyramidal neurons, GPSRs exerted inhibitory effects when uEPSPs were initiated at the GABA puff site (0 µm). However, unlike pyramidal neurons, GS GPSRs produced excitation for uEPSPs triggered at both distal (+100 µm) and proximal (−100 µm) dendritic locations (Fig. 2C, D; Supplementary Tables 5, 6). Although inhibition remained present at co-localized sites, its magnitude was significantly weaker than in pyramidal neurons (Supplementary Fig. 1). These findings suggest that the weaker shunting inhibition in GSs allows depolarizing GPSRs to exert excitatory effects even at distal dendritic locations (−100 µm).

**Figure 2.**
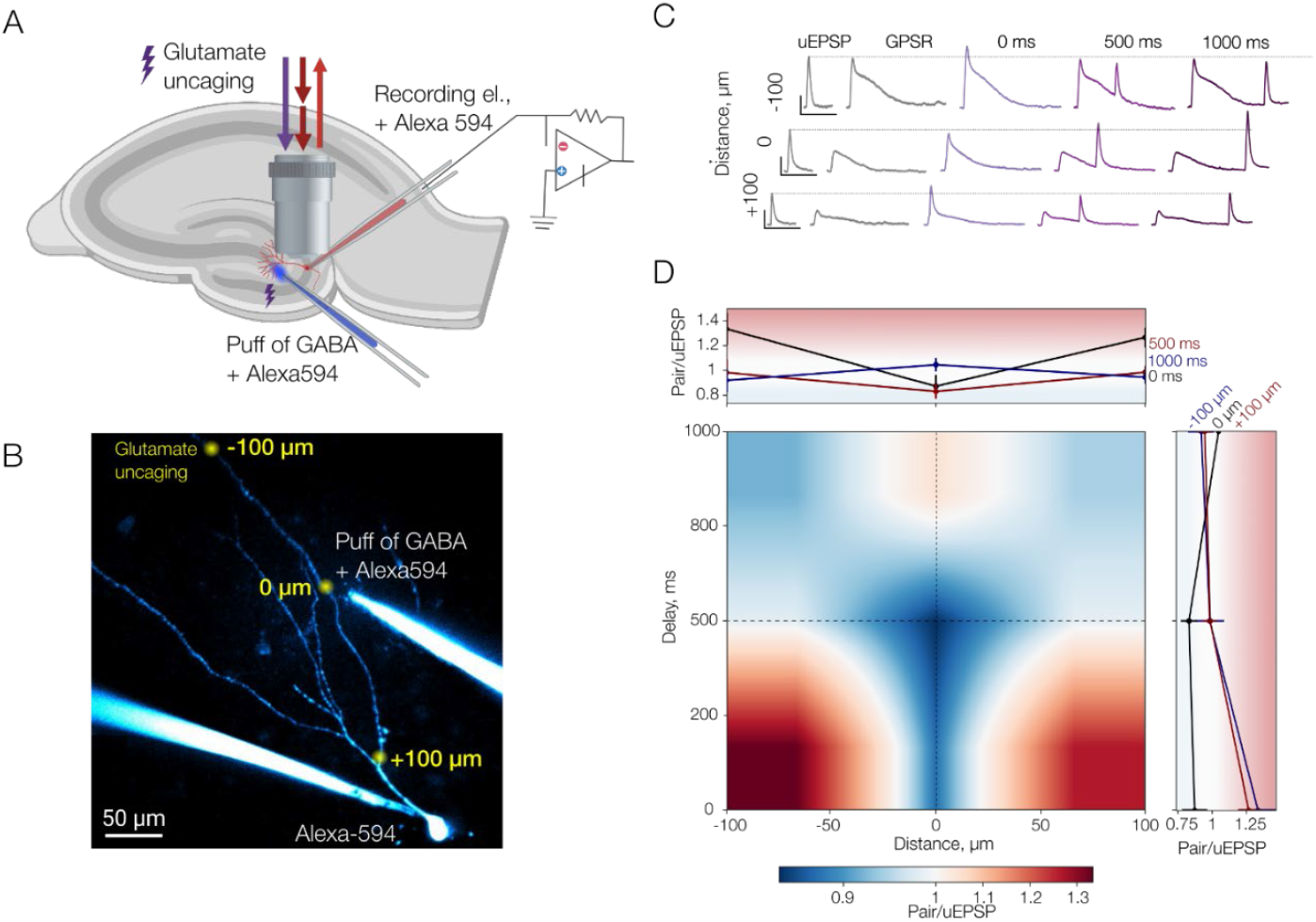
Depolarizing effect of GABA in dentate gyrus GSs. **A**. Schematic of hippocampal slice, recording electrode, and puff pipette located in dentate gyrus (DG). **B**. GS filled with Alexa Fluor 594 (50 μM) and puff pipette filled with Alexa Fluor 594 (15 μM). **C**. Sample traces of uEPSP, GPSR, and paired uEPSP + GPSR responses for depolarizing GABA. **D**. Summary graph showing Pair/uEPSP ratio for different time delays and distances for depolarizing GABA. The data are presented as the mean ± SEM. For statistical data, see Supplementary tables 5-6.

All recordings in the preceding experiments were obtained at the soma, the principal site of synaptic integration. However, somatic recordings cannot resolve local events at the dendritic site of EPSP initiation. EPSPs undergo transformation on the way to the soma — attenuated by electrotonic decay, modulated by dendritic voltage-gated conductances, and potentially reduced by intervening *en route* inhibition. Because patch-clamp recordings are not feasible at every dendritic location where uEPSPs were generated, we constructed a biophysically detailed, multi-compartmental model of a CA1 pyramidal neuron to investigate the spatial and temporal dynamics of GABAergic signaling inaccessible by direct experimentation (Fig. 3A).

**Figure 3.**
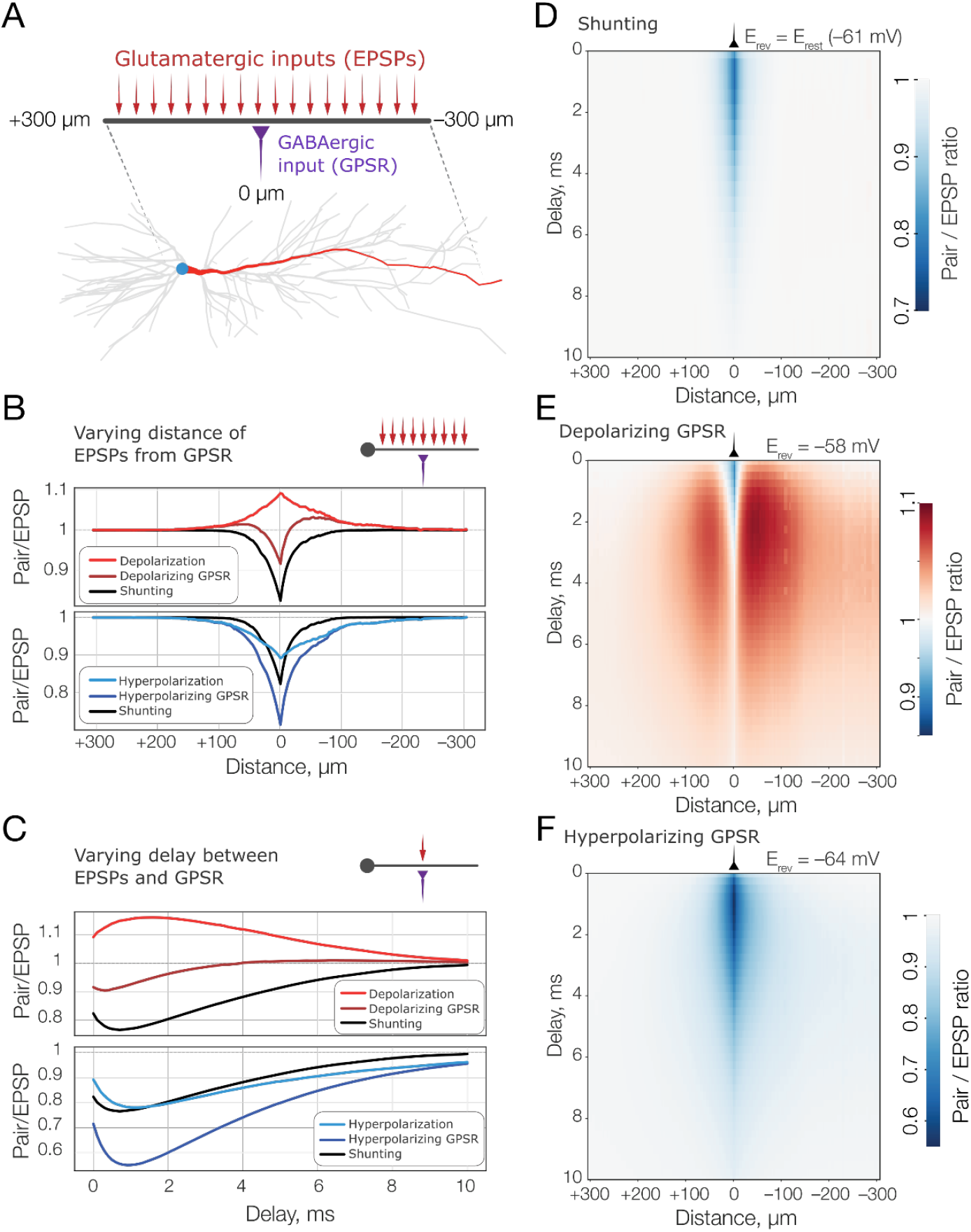
Simulation of depolarizing and hyperpolarizing effect of GABA in CA1 pyramidal neurons. **A**. Schematic of GPSR affecting EPSPs in a dendrite. **B**. Pair/uEPSP ratio at different distances for GPSR, pure shunting, pure polarization for depolarizing (*top*) and hyperpolarizing (*bottom*) GABA. **C**. Pair/uEPSP ratio for different delays for GPSR, pure shunting, pure polarization for depolarizing (*top*) and hyperpolarizing (*bottom*) GABA. **D**. Summary graph showing Pair/uEPSP ratio for different time delays and distances for pure shunting. **E**. Summary graph showing Pair/uEPSP ratio for different time delays and distances for depolarizing GABA. **F**. Summary graph showing Pair/uEPSP ratio for different time delays and distances for hyperpolarizing GABA.

The model incorporated a reconstructed CA1 pyramidal neuron morphology, with a GABAergic synapse fixed at the midpoint of the apical dendritic trunk (mirroring the experimental configuration) and a glutamatergic synapse movable along the trunk. Membrane potential was recorded at the site of glutamatergic input to quantify local EPSP amplitudes under the influence of GABAergic input.

We simulated three GABAergic reversal potentials to represent distinct modes of action: pure shunting (E_GABA_ = RMP = −61 mV), depolarizing (E_GABA_ = −58 mV), and hyperpolarizing (E_GABA_ = −64 mV). At each dendritic location, the glutamatergic conductance was calibrated to produce a 5-mV local EPSP in the absence of GPSR.

Consistent with experimental findings, both depolarizing and hyperpolarizing GPSRs reduced EPSP amplitude when colocalized with the glutamatergic input (distance 0 µm; Fig. 3B), with inhibition being stronger for hyperpolarizing GABA. When the glutamatergic input was spatially separated from the GABAergic input, inhibition declined. Notably, in the case of depolarizing GPSRs, the effect transitioned to excitation at 32.75 µm proximally and 16.57 µm distally relative to the GABAergic site—locations where depolarization overpowered the local shunting inhibition. In contrast, hyperpolarizing GPSRs remained inhibitory across all distances, although the effect weakened with increasing distance.

Unlike experimental approaches, the model enabled the separation of GPSR into its two constituent components: shunting and polarization. The pure shunting effect - defined as the effect when E_GABA_ = RMP - was identical for depolarizing and hyperpolarizing GPSRs. To isolate the polarization component, we subtracted the pure shunting response from the total response in both conditions. As expected, pure depolarization and hyperpolarization components exerted opposing effects, but both extended over broader spatial domains than the shunting component. Exponential fits revealed that the shunting effect decayed with a length constant of 23.25 µm, whereas polarization effects decayed with a longer length constant of 61.3 µm.

We also examined the distinct temporal dynamics of shunting and polarization (Fig. 3C). For co-localized GABAergic and glutamatergic synapses (0 µm), pure shunting inhibition peaked at 1 ms, reflecting the rapid kinetics of GABAA receptor-mediated conductance. In depolarizing GPSRs, shunting initially dominated, followed by a delayed depolarizing effect that resulted in weak excitation beginning 4 ms post-GABA_A_ receptor activation. Hyperpolarizing GPSRs exhibited sustained inhibition, shaped by the combined effects of shunting and hyperpolarization.

The integration of spatial and temporal dependencies yielded characteristic inhibitory– excitatory landscapes in two-dimensional parameter maps (Fig. 3D–F). Pure shunting inhibition was tightly confined in space and time (Fig. 3D). Depolarizing GPSRs produced complex, biphasic responses: inhibition at short distances and delays, followed by excitation as either distance or delay increased (Fig. 3E). Hyperpolarizing GPSRs induced robust inhibition throughout, with peak efficacy observed for spatially co-localized inputs at short delays (1 ms; Fig. 3F).

Together, these modeling results highlight the dual mechanisms of GABAergic inhibition, each with distinct spatiotemporal profiles. Shunting acts locally and transiently, whereas polarization spreads more broadly and persists longer. This interplay creates either biphasic (depolarizing) or synergistic (hyperpolarizing) effects on excitatory input. The net effect depends on synaptic positioning, timing, and dendritic biophysical properties, thereby defining dynamic integration windows shaped by the balance of membrane conductance and polarization.

## Discussion

The effects of GABAergic signaling on EPSPs are governed by the interplay between membrane polarization and conductance mediated by GABA_A_ receptor activation. Our study identifies three principal components underlying these effects: (1) shunting inhibition, which is spatially and temporally restricted; (2) membrane polarization, which spreads along the dendrite and outlasts the shunting component; and (3) *en route* inhibition, which occurs when an EPSP traverses a dendritic segment with active GABA_A_ receptors. While all three components scale with the number of activated GABA_A_ receptors, the polarization component is additionally determined by the GABA reversal potential (E_GABA_). Depolarizing GPSRs exhibited a biphasic effect - initial inhibition due to increased conductance, followed by delayed excitation driven by membrane depolarization. In contrast, hyperpolarizing GPSRs exerted consistently inhibitory effects.

Activation of GABAA receptors leads to a rapid increase in membrane conductance and the generation of local ionic currents that polarize the membrane (Takeuchi and Takeuchi, 1967). The temporal persistence of this polarization is governed by the membrane time constant, which reflects the ratio of membrane capacitance to conductance and varies across cell types (Gulledge and Stuart, 2003; Pugh and Jahr, 2011; Chiang et al., 2012; Haam et al., 2012). Once GABAergic currents subside, the membrane repolarizes for some time, allowing polarization to persist beyond the conductance phase (Lombardi et al., 2021b). This temporal dissociation enables depolarizing GABA to inhibit EPSPs during the conductance phase and subsequently excite the neuron via residual depolarization. Spatially, the distinction between the confined nature of shunting and the broader spread of polarization reflects a fundamental principle of dendritic integration. Importantly, this polarization is not strictly passive; activation of voltage-gated dendritic conductances may further shape and amplify the response (Song et al., 2011). Accordingly, changes in dendritic conductance - so-called dendritic plasticity - not only influence dendritic spike initiation (Losonczy et al., 2008; Gonzalez et al., 2022) but also modulate the integrative effects of GABAergic signaling.

The third component—*en route* inhibition—emerges when an EPSP propagates through a dendritic region with active GABAergic input. This mechanism can negate the excitatory impact of depolarizing GPSRs at distal dendritic sites, especially in CA1 pyramidal neurons, where we observed prominent *en route* inhibition. In contrast, this effect was absent or minimal in dentate gyrus GSs, potentially due to a weaker shunting component or enhanced EPSP amplification by dendritic conductances in these cells. Future investigations will be required to disentangle these possibilities.

The functional impact of GABAergic inputs also depends on their subcellular distribution. GABA_A_ receptors are differentially expressed across the soma, dendrites, and axon initial segment, each domain contributing uniquely to synaptic integration. Dendritic GABAergic synapses are well-positioned to locally modulate excitatory inputs (Kanemoto et al., 2011), while somatic synapses exert global control over output by influencing the summation of converging dendritic inputs. At the axon initial segment, GABAergic synapses precisely regulate AP generation by altering sodium channel availability and raising AP thresholds, thereby imposing spatially restricted shunting inhibition and delaying spike initiation (Lipkin and Bender, 2023). This subcellular compartmentalization underscores the specialized and context-dependent roles of GABA_A_ receptor-mediated inhibition.

The extent and duration of polarization are further shaped by membrane conductance, which is influenced by neuronal morphology and ion channel density. Neurons with larger surface area or higher channel expression have shorter membrane time and length constants, limiting the duration and spread of polarization. This may explain the differences observed in CA1 pyramidal neurons and dentate gyrus GSs (Kowalski et al., 2016). Developmental changes or pathological alterations in cell size or membrane properties - such as those occurring during pruning, growth, or neurodegeneration - could significantly impact GABAergic modulation of excitatory integration.

Another critical determinant is the developmental shift in E_GABA_ from depolarizing to hyperpolarizing, driven by alterations in chloride homeostasis via differential expression of KCC2 and NKCC1 transporters. This shift profoundly affects synaptic plasticity, switching the polarity of spike-timing-dependent plasticity (STDP) from Hebbian to anti-Hebbian in areas such as the striatum (Valtcheva et al., 2017). In pathological states, including epilepsy and traumatic brain injury, disruption of chloride regulation leads to depolarizing shifts in E_GABA_, which can impair inhibition and promote hyperexcitability (Watanabe and Fukuda, 2015; Hampel et al., 2021). Understanding the spatiotemporal components of GABAergic signaling - shunting, polarization, and *en route* inhibition - may offer therapeutic insights into restoring effective inhibition under such conditions.

In summary, our findings delineate the spatiotemporal dynamics of GABAergic and glutamatergic integration in the dendrites. The outcome of GABAA receptor activation is defined by the evolving interplay between membrane conductance and polarization, which together determine whether GABAergic inputs inhibit or excite, and under which conditions. These mechanisms differ across cell types, shaped by intrinsic membrane properties and synaptic organization. Through their influence on dendritic computation and network dynamics, GABAergic inputs play an essential role in both physiological processing and neurological disease states, including epilepsy (Feng et al., 2022; Perucca et al., 2023) and Alzheimer’s disease (Carello-Collar et al., 2023).

## Materials and Methods

### Hippocampal slice preparation

Transverse hippocampal slices were prepared from 3- to 5-week-old Sprague Dawley male rats in accordance with LASA ethical recommendations. Animals were anesthetized with 2-bromo-2-chloro-1,1,1-trifluroethane (halothane) and decapitated. The brain was exposed and cooled with an ice-cold solution containing (in mM): 75 sucrose, 87 NaCl, 2.5 KCl, 0.5 CaCl_2_, 1.25 NaH_2_PO_4_, 7 MgCl_2_, 25 NaHCO_3_, 1 Na-ascorbate, and 25 D-glucose. Hippocampi from both hemispheres were isolated and placed in an agar block. Transverse slices (350 µm) were prepared with a vibrating microtome (Microm HM 650V, Thermo Fisher Scientific Inc.) and left to recover for 20 min at 34°C and then for 40 min in an interface chamber with storage solution containing (in mM): 127 NaCl, 2.5 KCl, 1.25 NaH_2_PO_4_, 2 MgCl_2_, 1 CaCl_2_, 25 NaHCO_3_, and 25 D-glucose. The slices were then transferred to the recording chamber and continuously perfused with recording solution containing (in mM): 127 NaCl, 2.5 KCl, 1.25 NaH_2_PO_4_, 1 MgCl_2_, 2 CaCl_2_, 25 NaHCO_3_, and 11 D-glucose at 34°C. All solutions were saturated with 95% O_2_ and 5% CO_2_. Osmolarity was adjusted to 295 ± 5 mOsm; 5 mM 3-[[(3,4-dichlorophenyl) methyl]amino]propyl] diethoxymethyl) phosphonic acid (CGP52432) and 400 mM (S)-a-methyl-4-carboxyphenylglycine (S-MCPG) were routinely added to the solution to block GABA_B_ and metabotropic glutamate receptors, respectively. Cells were visually identified under infrared DIC using an Olympus BX-61 microscope (Olympus).

### Electrophysiology

Whole-cell recordings of the depolarizing effect of GABA in CA1 pyramidal neurons and GSs were obtained using patch electrodes filled with a solution containing (in mM): 125.67 CsCH_3_SO_3_, 4.43 KCl, 8 NaCl, 10 HEPES, 10 Na_2_-phosphocreatine, 4 Na_2_ATP, 0.4 NaGTP, 3 L-ascorbic acid. The solution was formulated for E_GABA_ = −62.81 mV (based on p_Cl_ : p_HCO2_ = 0.2 ((Kaila et al., 1997))), and the resting membrane potential = −67.07 mV (based on p_K_ : p_Na_ : p_Cl_ = 1 : 0.05 : 0.45 calculated by using the Goldman-Hodgkin-Katz equation). Whole-cell recordings of the hyperpolarizing effect of GABA in CA1 pyramidal neurons were obtained using patch electrodes filled with a solution containing (in mM): 130 CsCH_3_SO_3_, 5.7 NaCl, 10 HEPES, 11.15 Na_2_-phosphocreatine, 4 Na_2_ATP, 0.4 NaGTP, 3 L-ascorbic acid. The solution was formulated for E_GABA_ = −83.44 mV and the resting membrane potential = −72.66 mV. The pH of both solutions was adjusted to 7.2 with CsOH, and the osmolarity was adjusted to 290 mOsm. Pipette resistance was 3 – 5 MΩ. The recording solution also contained the morphological tracer Alexa Fluor 594 (50 μM) to visualize the soma and dendrites of the recorded neuron.

The signals were recorded with a patch-clamp amplifier Multiclamp 700B (Molecular Devices, USA), filtered at 4 kHz, and digitized at 10 kHz with an NI PCI-6221 card (National Instruments). The data were visualized and stored with the software WinWCP (supplied free of charge to academic users by Dr. John Dempster, University of Strathclyde, UK).

### Two-photon imaging

Cells were filled with the dye for at least 20 min before imaging to ensure subcellular dye distribution. Two-photon imaging was performed with a two-scanner FV1000-MPE laser-scanning microscope (Olympus) equipped with a mode-locked (<140 fs pulse width) tunable 720–930 nm laser Chameleon XR (Coherent, USA). Alexa Fluor 594 was excited at 830 nm light wavelength. We used the bright Alexa Fluor 594 emission to identify oblique apical dendrites (at 50-250 μm from the soma) and their spines to locally elicit Glutamate Uncaging-induced EPSCs (uEPSC) and GABAergic postsynaptic responses (GPSR = postsynaptic conductance + polarization)

### Glutamate Uncaging-induced EPSCs (uEPSC) and GABAergic postsynaptic responses (GPSR = postsynaptic conductance + polarization)

Bath application of 4-methoxy-7-nitroindolinyl-caged L-glutamate (MNI-glutamate; 400 μM) was used to ensure the stability of glutamate uncaging. Single-photon uncaging was carried out using 5–10 ms laser pulses (405 nm diode laser; FV5-LD405; Olympus) with ‘‘point scan’’ mode in Fluoview (Olympus). The uncaging spots were usually assigned at the edge of spine heads of imaged dendrites to elicit 5-7 mV uEPSC in soma. GABAergic postsynaptic responses (GPSRs) were triggered GABA puffs controlled with a picospritzer (General Valve, Fairfield, NJ) and delivered by a patch pipette at intervals of 15 s. The pipette was placed 50 –250 μm from the cell soma and contained GABA (10 μM) and Alexa Fluor 594F (15 μM) dissolved in (in mM): 127 NaCl, 2.5 KCl, 1.25 NaH_2_PO_4_, 25 NaHCO_3_, and 11 D-glucose; the pH was adjusted to 7.4 with NaOH. Delivery pressure was fixed at 6 psi, and the duration (8 –15 ms) was adjusted to yield 5-7 mV GPSRs in soma. Paired uEPSC and GPSR were induced at different points in time and space to measure how postsynaptic conductance + polarization affect uEPSP amplitude. We varied the time delay between uEPSC and GPSR (0, 200, 500, 800, and 1000 ms) and the spatial distance (−100, −50, 0, 50, and 100 μm) and measured the amplitude of uEPSP. To assess the spatiotemporal effect of GPSR on uEPSC, we measured the ratio between the response to paired stimulation and uEPSC stimulation alone.

### Drugs and chemicals

All drugs were made from stock solutions kept frozen at −20°C in 100 to 200 μl 1000x concentration aliquots. 3-[[(3,4-Dichlorophenyl)methyl]amin-o]propyl] diethoxymethyl) phosphinic acid (CGP 52432), (S)-α-Methyl-4-carboxyphenylglycine (S-MCPG), γ-Aminobutyric acid (GABA), D-(−)-2-Amino-5-phosphonopentanoic acid (APV), picrotoxin (PTX), 4-methoxy-7-nitroindolinyl-caged L-glutamate (MNI-glutamate) were purchased from Tocris Cookson (Bristol, UK). Alexa Fluor 594F was obtained from Invitrogen (Carlsbad, CA, USA).

### Data Analysis

Electrophysiological data were analyzed with WinWCP and Clampfit (Axon Instruments). Imaging data were visualized using FluoView (Olympus), ImageJ (a public domain Java image processing program by Wayne Rasband). Statistical analysis was performed using Excel (Microsoft, USA) and Origin 8 (OriginLab). The statistical significance was tested using a paired or unpaired Student’s *t*-test. The data are presented as mean ± SEM. ‘‘n’’ designates the number of recordings. In all figures, error bars indicate mean ± SEM.

### Neuron Model

To investigate signal integration across various locations on the dendritic tree, we developed a detailed biophysical simulation based on the CA1 pyramidal neuron model with reconstructed morphology by (Migliore et al., 2005). The model incorporated the following active conductances, with channel densities varying with distance from soma: voltage-gated sodium channels (Na_v_), delayed rectifier potassium channels (K_DR_), A-type potassium channels (K_A_ with proximal and distal variants), and H-channels (I_h_).

All parameters for active conductances and channel densities were the same as in (Migliore et al., 2005). The passive membrane properties were membrane capacitance C_m_ = 1 µF/cm2, axial resistance Ra = 150 Ω · cm for soma and dendrites and 50 Ω · cm for axon, and membrane resistance R_m_ = 28000 Ω · cm^2^. The reversal potentials were set to E_Na_ = 55 mV, E_K_ = −90 mV, Eh = −30 mV, and E_leak_ = −65 mV. All simulations were performed at the temperature of 35°C using the CVODE integration method.

### Synapse model

We studied the spatiotemporal interactions between GABAergic and glutamatergic synapses by placing a pair of synapses at variable locations along the shaft. Synapses were modeled using the Exp2Syn mechanism in NEURON, which implements a two-state kinetic scheme with exponential rise and decay:

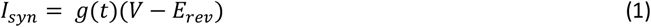

where the time-dependent conductance g(t) is given by:

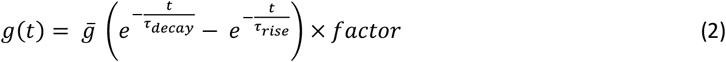

Here:

*I*_*syn*_ - synaptic current, *g*(*t*)-time-dependent conductance, *V* - membrane potential, *E*_*rev*_ - synaptic reversal potential, 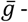 peak synaptic conductance, *τ*_*rise*_, *τ*_*decay*_ - rise and decay time constants, 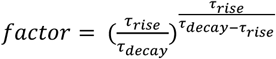, normalizing the conductance so that 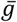 represents the peak conductance (for parameters, see Table 1).

**Table 1:** Parameters for glutamatergic and GABAergic synapses used in the model.

|  | Glu synapse | GABA synapse |
| --- | --- | --- |
| $\tau_{rise}$ | 0.1 ms | 0.5 ms |
| $\tau_{decay}$ | 1 ms | 3 ms |
| $E_{rev}$ | 0 mV | variable |
| $\bar{g}$ | variable | 0.50 nS |

### Conductance calibration

Because the original model contained active dendritic conductance, strong inputs could trigger dendritic spikes. To avoid dendritic spike generation, we performed a calibration of glutamatergic synaptic conductances 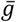Glu. At each location along the dendritic shaft, we adjusted the value of 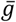Glu to elicit a subthreshold depolarization of 5 mV at the synapse. The calibration was performed using a binary search algorithm in the range from 0 to 0.1 µS.

### Simulation protocol

After calibrating glutamatergic conductances, we introduced a GABAergic synapse at the midpoint of the apical dendrite. To examine different modes of GABAergic signaling, we tested three E_GABA_: 1. Shunting: E_GABA_ = RMP = −60.61 mV; 2. Depolarizing GABA: E_GABA_ = −58 mV; 3. Hyperpolarizing GABA: E_GABA_ = −64 mV.

To explore spatiotemporal interactions between GPSR and EPSP, we varied the location of the glutamatergic synapse along the dendritic shaft and the temporal delay between GABAergic and glutamatergic synapse activation. The temporal delays ranged from 0 to 10 ms in 0.25 ms increments. Each simulation ran for 100 ms with a time step of 0.025 ms. Before synaptic activation, the model was allowed to settle into the equilibrium for 100 ms.

For each simulation, we recorded the membrane potential at the glutamatergic synapse location to analyze local EPSP. We quantified the GPSR effect by calculating the ratio between peak EPSP amplitudes with and without GPSR. While our standard protocol used temporal delays from 0 to 10 ms in 0.25 ms increments, we performed additional high-resolution simulations for colocalized GABA and glutamate synapses at the shaft midpoint using smaller temporal delays of 0.1 ms to better capture the effect.

## Supplementary materials

**Supplementary video 1. Puff of GABA onto a dendrite of CA1 pyramidal neurons**. Two-photon imaging of CA1 pyramidal neuron filled with Alexa Fluor 594 (50 μM). A glass puff pipette was placed in CA1 stratum radiatum next to a dendrite and contained GABA (10 μM) and Alexa Fluor 594F (15 μM).

**Supplementary Fig. 1.**
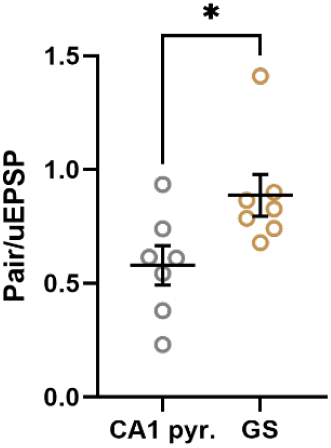
The inhibitory effect of depolarizing GABA is weaker in GS than in CA1 pyramidal neurons. Summary graph showing Pair/uEPSP ratios in CA1 pyramidal neurons and dentate gyrus GS for colocalized GPSR and uEPSP at zero delay. The data are presented as the mean ± SEM. Circles represent individual recordings. * - p < 0.05, two-sided *t*-test

**Supplementary Table 1.** The effect of depolarizing GPSR on uEPSP amplitude in CA1 pyramidal neuron, n = 7, EPSP/Pair ± SEM, one sample t-test for comparison to 1, p-value, * - p < 0.05, ** - p < 0.01.

| Distance ($\mu\text{m}$) / Delay (ms) | 0 ms | 200 ms | 500 ms | 800 ms | 1000 ms |
| --- | --- | --- | --- | --- | --- |
| 100 $\mu\text{m}$ | 1.06 ± 0.06, 0.3404 | 1.20 ± 0.07, 0.0268, * | 1.21 ± 0.04, 0.0031, ** | 1.06 ± 0.04, 0.1797 | 1.05 ± 0.03, 0.1706 |
| 50 $\mu\text{m}$ | 0.69 ± 0.12, 0.0422, * | 0.94 ± 0.08, 0.471 | 1.00 ± 0.07, 0.9517 | 0.99 ± 0.04, 0.8177 | 1.11 ± 0.02, 0.0043, ** |
| 0 $\mu\text{m}$ | 0.63 ± 0.11, 0.0146, * | 0.74 ± 0.07, 0.0115, * | 0.82 ± 0.07, 0.0453, * | 0.91 ± 0.06, 0.2162 | 0.98 ± 0.03, 0.4759 |
| -50 $\mu\text{m}$ | 0.77 ± 0.12, 0.0621 | 0.82 ± 0.10, 0.0716 | 0.85 ± 0.08, 0.1826 | 0.88 ± 0.06, 0.2168 | 0.93 ± 0.06, 0.3980 |
| -100 $\mu\text{m}$ | 0.77 ± 0.12, 0.1242 | 0.96 ± 0.11, 0.7032 | 0.95 ± 0.05, 0.3956 | 0.97 ± 0.03, 0.4409 | 0.95 ± 0.07, 0.5047 |

**Supplementary Table 2.** Spatial-temporal effect of depolarizing GPSR on uEPSP amplitude in CA1 pyramidal neurons, n = 7, Two-way RM ANOVA, * - p < 0.05, *** - p < 0.001.

| Spatial condition | F, df, P value | Temporal condition | F, df, P value |
| --- | --- | --- | --- |
| 0 $\mu\text{m}$ vs. 100 $\mu\text{m}$ | F(1, 48) = 46.62, p<0.0001*** | 0 ms vs. 200 ms | F(1, 48) = 4.76, p=0.034* |
| 0 $\mu\text{m}$ vs. 50 $\mu\text{m}$ | F(1, 48) = 11.71, p=0.0013 * | 0 ms vs. 500 ms | F(1, 48) = 9.56, p<0.01** |
| 0 $\mu\text{m}$ vs. -50 $\mu\text{m}$ | F(1, 48) = 0.44, p=0.5068 | 0 ms vs. 800 ms | F(1, 48) = 14.86, p<0.0001*** |
| 0 $\mu\text{m}$ vs. -100 $\mu\text{m}$ | F(1, 48) = 2.29, p=0.1362 | 0 ms vs. 1000 ms | F(1, 48) = 23.43, p<0.0001*** |

**Supplementary Table 3.** The effect of hyperpolarizing GPSR on uEPSP amplitude in CA1 pyramidal neuron, n=6, EPSP/Pair ± SEM, one sample t-test for comparison to 1, p-value, * - p<0.05, ** - p<0.01, *** - p<0.001.

| Distance ($\mu\text{m}$) / Delay (ms) | 0 ms | 200 ms | 500 ms | 800 ms | 1000 ms |
| --- | --- | --- | --- | --- | --- |
| 100 $\mu\text{m}$ | 0.43 ± 0.07, 0.0003, *** | 0.29 ± 0.11, 0.0019, *** | 0.46 ± 0.11, <0.0001, *** | 0.60 ± 0.12, <0.0001, *** | 0.74 ± 0.09, 0.006, ** |
| 50 $\mu\text{m}$ | 0.30 ± 0.12, 0.0012, ** | 0.05 ± 0.09, 0.0001, *** | 0.29 ± 0.07, <0.0001, *** | 0.52 ± 0.09, <0.0001, *** | 0.65 ± 0.12, <0.0001, *** |
| 0 $\mu\text{m}$ | -0.33 ± 0.10, 0.0044, ** | -0.16 ± 0.05, 0.0001, *** | 0.13 ± 0.08, 0.0002, *** | 0.54 ± 0.14, 0.0014, ** | 0.70 ± 0.14, 0.0006, *** |
| -50 $\mu\text{m}$ | -0.33 ± 0.04, 0.0197, * | -0.15 ± 0.04, 0.0036, ** | 0.04 ± 0.15, 0.0192, * | 0.58 ± 0.13, 0.0235, * | 0.71 ± 0.14, 0.0225, * |
| -100 $\mu\text{m}$ | -0.18 ± 0.26, 0.0327, * | -0.10 ± 0.09, 0.033, * | 0.18 ± 0.11, 0.0812 | 0.56 ± 0.14, 0.0899 | 0.66 ± 0.13, 0.0471, * |

**Supplementary Table 4.** Spatial-temporal effect of hyperpolarizing GPSR on uEPSP amplitude in CA1 pyramidal neurons, n = 6, Two-way RM ANOVA, * - p < 0.05, ** - p < 0.01, *** - p < 0.001.

| Spatial condition | F, df, P value | Temporal condition | F, df, P value |
| --- | --- | --- | --- |
| 0 $\mu\text{m}$ vs. 100 $\mu\text{m}$ | F(1, 40) = 8.31, p=0.0063** | 0 ms vs. 200 ms | F(1, 40) = 0.28, p= 0.5999 |
| 0 $\mu\text{m}$ vs. 50 $\mu\text{m}$ | F(1, 40) = 7.95, p=0.0069** | 0 ms vs. 500 ms | F(1, 40) = 9.85, p= 0.0032* |
| 0 $\mu\text{m}$ vs. -50 $\mu\text{m}$ | $F(1, 40) = 0.01, p = 0.9305$ | 0 ms vs. 800 ms | $F(1, 40) = 48.1, p < 0.0001^{***}$ |
| 0 $\mu\text{m}$ vs. -100 $\mu\text{m}$ | $F(1, 40) = 0.08, p = 0.7776$ | 0 ms vs. 1000 ms | $F(1, 40) = 65.41, p < 0.0001^{***}$ |

**Supplementary Table 5.** The effect of depolarizing GPSR on uEPSP amplitude in granule cells in dentate gyrus, n = 6, EPSP/Pair ± SEM, one sample t-test for comparison to 1, p value, * - p < 0.05.

| Time point ($\mu\text{m}$) / Delay (ms) | 0 ms | 500 ms | 1000 ms |
| --- | --- | --- | --- |
| 100 $\mu\text{m}$ | $1.27 \pm 0.08, 0.0208, *$ | $0.98 \pm 0.05, 0.7875$ | $0.95 \pm 0.05, 0.3527$ |
| 0 $\mu\text{m}$ | $0.87 \pm 0.09, 0.2166$ | $0.83 \pm 0.06, 0.0299, *$ | $1.05 \pm 0.05, 0.4322$ |
| 100 $\mu\text{m}$ | $1.33 \pm 0.13, 0.0479, *$ | $0.98 \pm 0.11, 0.8734$ | $0.92 \pm 0.10, 0.4478$ |

**Supplementary Table 6.** Spatial-temporal effect of depolarizing GPSR on uEPSP amplitude in granule cells in dentate gyrus, n = 6, Two-way RM ANOVA

| Spatial condition | F, df, P value | Temporal condition | F, df, P value |
| --- | --- | --- | --- |
| 0 $\mu\text{m}$ vs. 100 $\mu\text{m}$ | $F(1, 20) = 0.01, p = 0.9005$ | 0 ms vs. 200 ms | $F(1, 20) = 3.29, p = 0.0845$ |
| 0 $\mu\text{m}$ vs. -100 $\mu\text{m}$ | $F(1, 20) = 0.01, p = 0.9395$ | 0 ms vs. 1000 ms | $F(1, 20) = 1.21, p = 0.2841$ |

